# Metastable GPCR dimers trigger the basal signal by recruiting G-proteins

**DOI:** 10.1101/2020.02.10.929588

**Authors:** Rinshi S. Kasai, Takahiro K. Fujiwara, Akihiro Kusumi

## Abstract

G-protein-coupled receptors (GPCRs) constitute the largest family of integral membrane proteins in the human genome and are responsible for various important signaling pathways for vision, olfaction, gustation, emotion, cell migration, etc. A distinct feature of the GPCR-family proteins is that many GPCRs, including the prototypical GPCR, β2-adrenergic receptor (β2AR), elicit low levels of basal constitutive signals without agonist stimulation, which function in normal development and various diseases^1–3^. However, how the basal signals are induced is hardly known. Another general distinctive feature of GPCRs is to form metastable homo-dimers, with lifetimes on the order of 0.1 s, even in the resting state. Here, our single-molecule-based quantification^4^ determined the dissociation constant of β2AR homo-dimers in the PM (1.6 ± 0.29 copies/μm^2^) and their lifetimes (83.2 ± 6.4 ms), and furthermore found that, in the resting state, trimeric G-proteins were recruited to both β2AR monomers and homo-dimers. Importantly, inverse agonists, which suppress the GPCR’s basal constitutive activity, specifically blocked the G-protein recruitment to GPCR homo-dimers, without affecting that to monomers. These results indicate that the G-proteins recruited to transient GPCR homo-dimers are responsible for inducing their basic constitutive signals. These results suggest novel drug development strategies to enhance or suppress GPCR homo-dimer formation.

## β2AR forms metastable homo-dimers

Single-molecule imaging-tracking revealed that β2AR (tagged with ACP at the N-terminus and labelled with ATTO594; **Extended Data Fig. 1**) undergoes thermal diffusion in the plasma membrane (PM), and often forms transient homo-dimers, undergoing rapid interconversions between monomers and dimers continually, as found for other GPCRs (**Fig. 1a, Extended Data Fig. 2a**)^4–7^. Every time we found a β2AR dimer, we measured the dimer duration, and after observing many dimers, we obtained the distribution of dimer durations (**Fig. 1b, Extended Data Fig. 2d**). The histogram could be fitted by a single exponential function, providing the exponential dimer lifetime of 99 ms (throughout this report, all statistical parameters, including SEM, and the number of experiments are summarised in **Extended Data Tables 1-4**; all observations were performed at 37°C). The lifetime value obtained in this manner, termed *τ*_observed_, was corrected for the photobleaching lifetime and incidental colocalisation lifetime (this method was used for evaluating all of the lifetimes investigated in the present research; see **Methods**), giving a dimer lifetime *τ* of 83.2 ± 6.4 ms (**Fig. 1c**). The dimer lifetime obtained at a time resolution of 4 ms (250 Hz) was 78.8 ± 10.7 ms, after corrections for photobleaching and incidental colocalisation lifetime (**Extended Data Fig. 3, Fig. 1c**), which is statistically indistinguishable from the dimer lifetime of 83.2 ms evaluated from 30-Hz observations. This result indicates that a frame rate of 30 Hz is sufficiently fast for evaluating the homo-dimer lifetime of β2AR. Together, these results demonstrate that the β2AR homo-dimers are indeed quite metastable, turning into monomers in a matter of 80 ms, and then quite readily forming another dimer with a different partner molecule (**Extended Data Fig. 2a**).

**Fig. 1.**
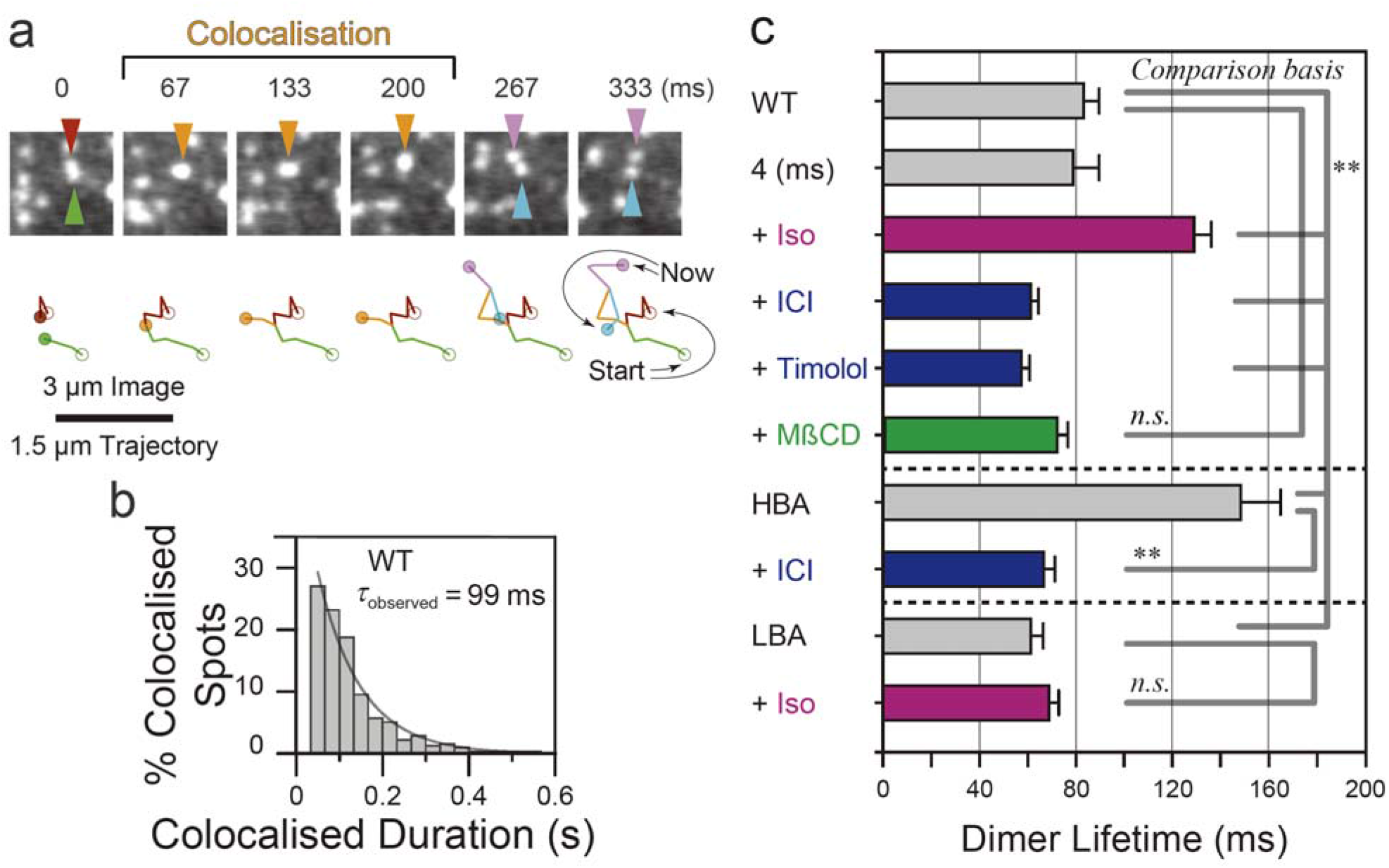
β2AR dynamically interconverts between monomers and transient homodimers. Mean, SEM, the number of experiments, and other statistical parameters are summarised in **Extended Data Table 1**. **(a)** A typical snapshot sequence showing transient homo-dimerisation of β2AR molecules. Two molecules became colocalised at frame 2 (67 ms), diffused together for 133 ms (colocalisation), and then separated. **(b)** Histogram showing the distribution of the durations of the WT-β2AR colocalisation events (homo-dimers). Note that the *τ*_observed_ is the value before corrections for photobleaching and incidental colocalisations. **(c)** Summary of the dimer lifetimes (after corrections) of WT and mutant β2AR obtained before and after the additions of various reagents. ** and *n.s*.: statistically significant and non-significant with *P* values smaller or greater than 0.05, respectively. This convention is used throughout this report.

## 2D-dissociation constant for β2AR dimers

Using single-molecule-based “super-quantification” method previously developed by us^4^, we obtained the 2-dimensional dissociation constant (2D-*K*_D_) of β2AR homo-dimers in the PM of living cells, which was 1.6 ± 0.29 copies/μm^2^ (**Fig. 2a, Extended Data Fig. 2b**; see **Methods**). This value is comparable to the 2D-*K*_D_ of another GPCR, formyl-peptide receptor (FPR), which was 3.6 copies/μm^2^ (in fact, this was the first determination of the 2D-*K*_D_ in the PM of living cells)^4^.

**Fig. 2.**
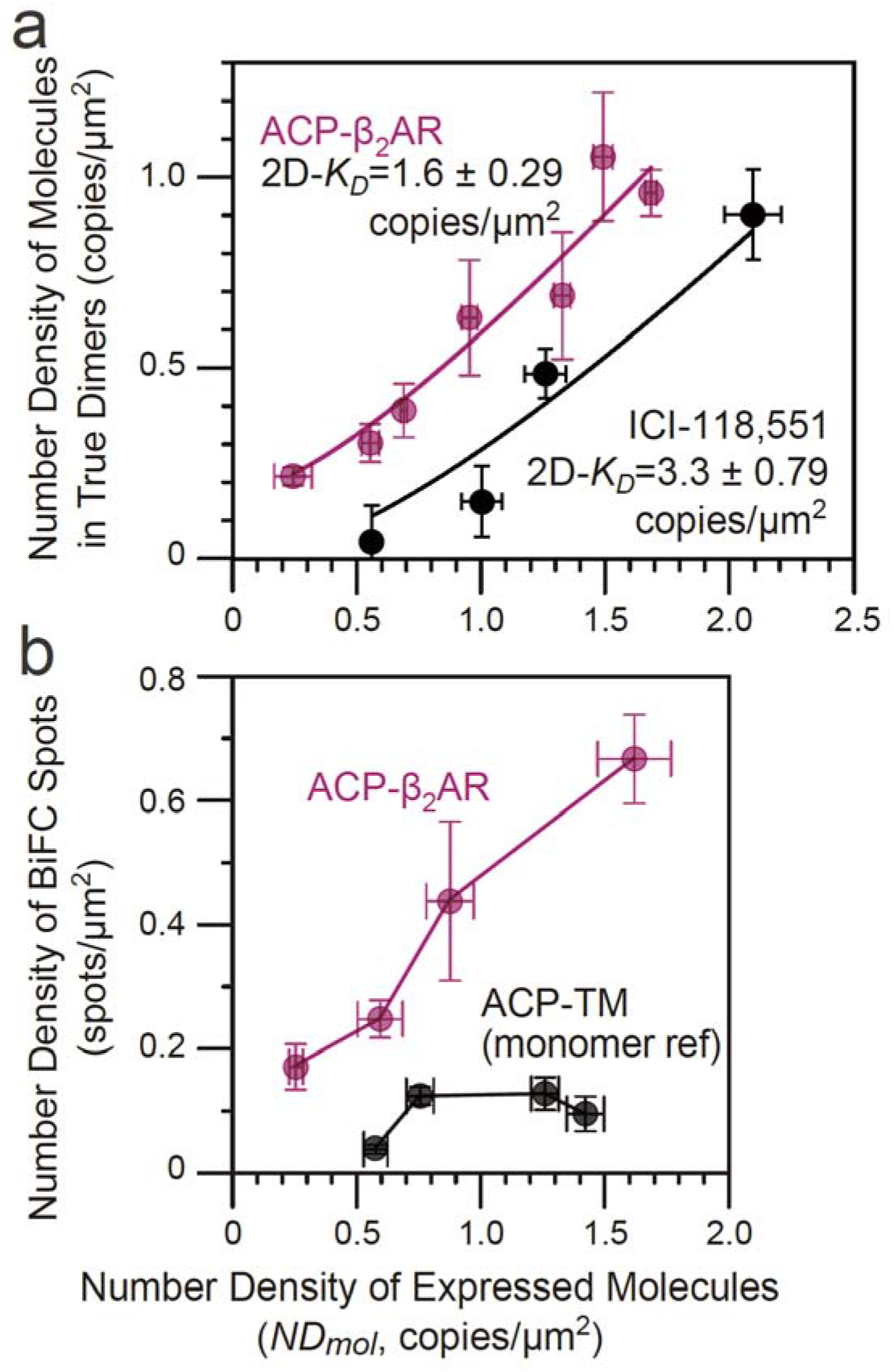
Determining the 2D-*K*D of β2AR dimers with/without an inverse agonist and the BiFC detection of β2AR dimers. Mean, SEM, the number of experiments, and other statistical parameters are summarised in **Extended Data Table 2**. The number density of β2AR molecules in true dimers (after subtraction of incidental overlap of fluorescent spots; ± inverse agonist ICI-118,551; curves represent the fitting results using Eq. 5 in Ref. 4) **(a)** and the number densities of the BiFC spots for ACP-β2AR-YFP(N/C) and ACP-TM-YFP(N/C) (monomer reference molecule) **(b)**, plotted as a function of the number density of expressed molecules (*NDmol*). In **b**, the lines are provided to facilitate visualisation.

Under physiological conditions, β2AR is expressed at number densities of 16 ~ 260 copies/μm^2^, depending on the cell type^8,9^. From the 2D-*K*_D_ determined here, we conclude that 12 ~ 240 copies/μm^2^ exist as dimers and 4 ~ 20 copies/μm^2^ exist as monomers at any moment; *i.e*., 75 ~ 92% of β2AR molecules exist as homo-dimers at any moment, although dimers and monomers are continually interconverting rapidly all the time (with a dimer lifetime of ~80 ms). In contrast, FPR is mainly expressed in neutrophils, and an average of about 6,000 copies of FPR copies exist in the PM of a single neutrophil^10^. Assuming that the neutrophil surface area is 1,300 μm^2^ (approximated by a sphere with a radius of 10 μm, providing a number density of ~4.6 FPR copies/μm^2^) and based on the 2D-*K*_D_ of FPR (3.6 copies/μm^2^), 3,200 copies of FPR would exist as monomers and 2,800 copies as dimers; namely, 54% of FPR molecules are dimers, which is about two-thirds of the dimer fraction of β2AR. Note that, even under the conditions where the dimer population dominates (*i.e*., at higher expression levels), the dimer lifetimes do not change and remain on the order of 0.1 s. Only the monomer lifetimes are shortened at higher expression levels.

## Direct detection of β2AR dimers by the BiFC method

Thus far, we have detected dimers by the colocalisation of two single molecules, which reveals the close encounters of molecules in the space scale of 220 nm, a distance much greater than the molecular scale. Therefore, in this method, the incidental approaches of two molecules were evaluated and subtracted. To ascertain the molecular-level interactions (dimers), we employed YFP-based bimolecular fluorescence complementation (BiFC; meanwhile, our attempt to detect FRET did not work, perhaps because the distance between the probes attached to two C-termini was too far for FRET)^4,11^. In BiFC, two potentially interacting proteins are fused to the N- and C-terminal half-molecules of YFP. If the proteins of interest form dimers, then YFP may be reconstituted and emit fluorescence signals (see **Methods**). Thus, if BiFC occurs, it would lend strong support for the formation of β2AR dimers.

To quantitatively examine BiFC, we varied the expression levels of β2AR. With an increase in the number density of expressed molecules (determined by the signal from ATTO594 bound to ACP; the ATTO594/ACP ratio is >0.95^12^), the number density of the BiFC spots of β2AR increased, whereas the BiFC-spot number density of the monomer reference molecule, ACP-TM, increased only slightly (**Fig. 2b; Extended Data Fig. 2c**). This result indicates that β2AR forms directly-bound dimers.

## Inverse agonists reduce the β2AR dimer affinity

Treatments with the inverse agonists ICI-118,551 (ICI) and timolol reduced the β2AR dimer lifetime (0.74x and 0.69x, respectively; **Fig. 1c, Extended Data Fig. 2d**), whereas ICI increased the 2D-*K*_D_ of β2AR by a factor of 2.1 (lowered the β2AR dimer affinity; **Fig. 1c**), indicating that inverse agonists reduce the β2AR dimer affinity. Meanwhile, the addition of the agonist isoproterenol (Iso) prolonged the dimer lifetime significantly (1.55x) (**Fig. 1c; Extended Data Fig. 2d**). Cholesterol reportedly modulates the functions of some GPCRs^13^, but here we found that mild cholesterol depletion by methyl-β-cyclodextrin (MβCD) treatment failed to affect the β2AR dimer lifetime (**Fig. 1c**).

Since the inverse agonists suppressed the basal constitutive GPCR signaling, and since we found that they also reduced the β2AR dimer lifetime, we next examined the dimer lifetimes of the β2AR mutants with high and low basal activities in the resting state (HBA and LBA mutants, representing E268A and C327R mutants, respectively^14,15^). The HBA mutant exhibited a 1.78x longer dimer lifetime, and interestingly, after the inverse-agonist addition, it was reduced to the dimer lifetime of the wild-type receptor (**Fig. 1c; Extended Data Fig. 2d**). Meanwhile, the LBA mutant exhibited a 0.74x shorter dimer lifetime than that of the wild-type receptor, and the agonist addition did not affect it (**Fig. 1c; Extended Data Fig. 2d**).

Strikingly, the dimer lifetimes of WT-β2AR after the addition of inverse agonists were quite comparable to those of the LBA mutant, and the dimer lifetime of WT-β2AR after the addition of the agonist isoproterenol was comparable to that of the HBA mutant. These results suggest that the β2AR conformations would be quite different after the binding of the inverse agonists and the agonist isoproterenol, as compared with those before their additions^16^, possibly resembling the conformations of the LBA and HBA mutants, respectively, and detectable as changes in the homo-dimer lifetimes.

## The basal constitutive activity of β2AR

We confirmed the basal constitutive activity of β2AR, as well as the effects of the β2AR agonist isoproterenol and the inverse agonists ICI-118,551 and timolol, by measuring the changes in the cytoplasmic cAMP levels in the L-cell line stably expressing β2AR at higher levels (**Extended Data Fig. 4, Extended Data Table 4**). The addition of the agonist isoproterenol to the cells increased the cAMP level by a factor of ~4, relative to the control. Meanwhile, the addition of the inverse agonists ICI-118,551 and timolol reduced the cAMP levels by factors of 1.9 and 1.3 (53 and 76% of the control), respectively, showing the presence of the basal constitutive activity of β2AR and the effectiveness of the inverse agonists, ICI-118,551 and timolol, for strongly reducing this basal activity.

## Trimeric G proteins are recruited to both β2AR monomers and dimers

The presence of the basal constitutive activity of β2AR suggests that a stimulatory trimeric G-protein might be recruited to β2AR in the resting state^17^. Therefore, we examined whether Gαs and Gβ(conjugated with mGFP) are recruited to β2AR monomers or dimers, by examining their colocalisations. Using L cells expressing both β2AR (ATTO594-ACP-β2AR) and Gαs or Gβ, we found that, even in resting cells, Gαs and Gβ were recruited to both β2AR monomers and dimers (**Fig. 3a; Supplementary Videos 1 and 2**).

**Fig. 3.**
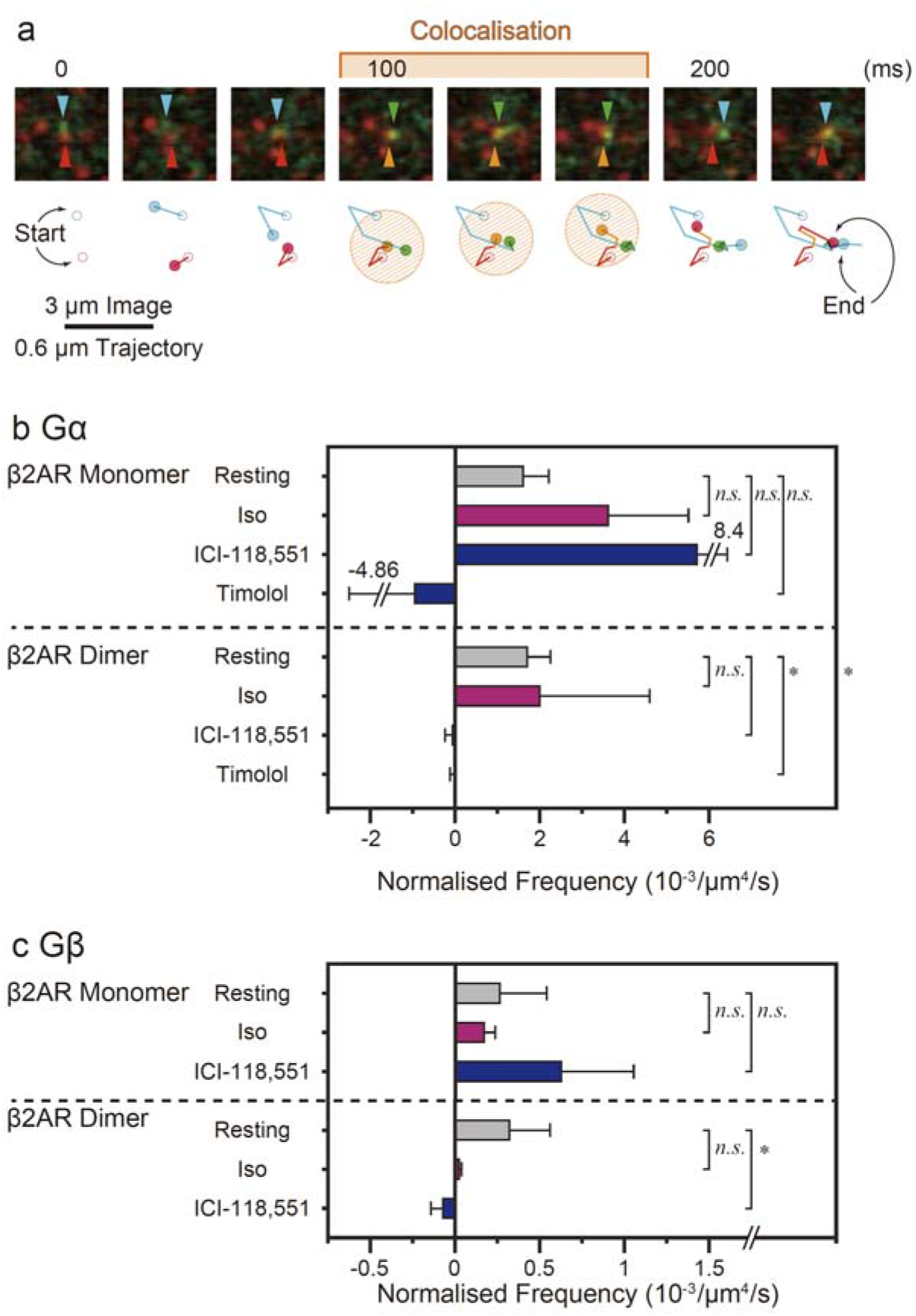
Non-stimulated β2AR homo-dimers recruit trimeric G-proteins: a process suppressed by inverse agonists. Mean, SEM, the number of experiments, and other statistical parameters are summarised in **Extended Data Table 3**. **(a)** A typical snapshot image sequence showing that a trimeric G-protein Gαs subunit transiently binds to a β2AR monomer. **(b, c)** The inverse agonists, ICI-118,551 and timolol, inhibitors of β2AR basal constitutive activities, blocked the recruitment of trimeric G-proteins to β2AR homo-dimers, but not that to monomers. Gαs **(b)** and Gβγ **(c)** colocalisation frequencies with β2AR monomers and homo-dimers, normalised by the number densities of Gαs (or Gβγ) and β2AR protomers (per sec per μm^4^).

The recruitment frequencies of Gαs and Gβ to β2AR monomers and dimers were measured and then normalised by the number densities of Gαs (or Gβγ) and β2AR protomers in the PM (normalised frequencies, with a dimension of per sec per μm^4^; **Fig. 3b, c; Extended Data Table 3**)^18^. In resting cells, both Gαs and Gβmolecules were recruited to β2AR monomers or dimers equally well, suggesting that (non-stimulated) β2AR monomers and/or dimers might induce intracellular Gs-dependent signals even in the resting state, which would lead to the basal constitutive activity of β2AR.

The exponential dwell lifetimes of G proteins on β2AR were on the order of 50 ~ 60 ms for all of the combinations of Gαs and Gβ molecules vs β2AR monomers and dimers (**Extended Data Fig. 5**). However, these durations are only the maximal estimates of the dwell lifetimes of Gαs and Gβ molecules on β2AR monomers and dimers, due to the present instrumental limitations of detecting short binding durations.

Notably, the treatment of the cells with the agonist, isoproterenol, did not change the recruitment frequencies in any combinations of Gαs and Gβ vs β2AR monomers and dimers, although it significantly enhanced the cytoplasmic cAMP level (**Extended Data Fig. 4, Extended Data Table 4**). This result is quite surprising, but it suggests that the isoproterenol-induced β2AR conformation enhanced its GEF activity, rather than increasing the on-rate of G protein binding to β2AR.

## Inverse agonists specifically block G-protein recruitment to β2AR dimers

Next, we examined the effects of the inverse agonists. The recruitment of Gαs and Gβ to β2AR dimers was almost completely blocked, whereas that to β2AR monomers was unaffected (**Fig. 3b, c**). This result unequivocally shows that β2AR dimers are responsible for the basal constitutive activity of β2AR. Furthermore, it demonstrates that the inverse agonists work by blocking the recruitment of stimulatory G-proteins to β2AR dimers, perhaps by inducing β2AR conformational changes in the binding sites for G-proteins, rather than blocking the β2AR dimerisation (although, as already described, the dimer affinity of the inverse agonist-bound β2AR is lower, resembling the LBA mutant, and thus inverse agonists might change the GPCR conformations in two ways: lowering the GPCR dimer affinity and modulating the G-protein binding sites).

To further confirm that β2AR dimers trigger downstream signaling without an agonist, and to detect the basal dimer signal at very low β2AR expression levels used for single-molecule experiments (generally <1 copy/μm^2^, which is approximately comparable to <3,000 copies in the PM/cell), we examined whether artificially-induced stable β2AR dimers could elicit the downstream signal. To detect the downstream signal at very low expression levels of β2AR, which are necessary for detecting dimers by single-molecule imaging, and also to detect the signal in a different manner, rather than observing the cAMP level again (**Extended Data Fig. 4**), we examined the intracellular Ca^2+^ mobilisation, which could occur further downstream from the cAMP amplification^19^. The induction of the artificially-induced stable β2AR (conjugated to FKBP) dimers by the FKBP-A20187 dimerisation system indeed elicited Ca^2+^ mobilisation (**Extended Data Fig. 6**). This result further confirmed that β2AR dimers could trigger the downstream signals without agonist stimulation.

## Discussion

All of the class-A GPCRs examined thus far, including β2AR examined here and by others^5^, FPR, D2 dopamine receptor, M1 muscarinic receptor, and α1AR, form metastable dimers with lifetimes on the order of 0.1 s in the non-stimulated state^4–7^. Therefore, we propose that the dynamic equilibrium between monomers and homo-dimers is a common characteristic property of the class-A GPCRs. The homo-dimerisation of GPCRs without any noticeable amino-acid homologies suggests that, through evolution, GPCRs changed almost all of their amino acids, but conserved the physical ability to form transient homo-dimers. This infers that these dynamic dimers are critical for some of the GPCR functions.

GPCRs are distinct among other receptors, in that, under non-stimulated conditions, GPCRs maintain the low basal constitutive activity, which can be inhibited by inverse agonists. The basal activities of GPCRs are implied in normal development and various diseases^1–3^. Therefore, we examined the possibility that the function of the GPCR dimers is to induce the basal constitutive activity in resting cells (without agonists), and obtained three unexpected, surprising findings. First, both β2AR monomers and dimers continually recruit G proteins, one molecule after another, even in the resting state. Second, the G-protein recruitment frequencies in the resting state are about the same in the presence of the agonist. Third, more importantly, the inverse agonists specifically and almost completely blocked the recruitment of Gαs and Gβ to β2AR dimers, without affecting their recruitment frequencies to β2AR monomers (**Fig. 3b, c**). The third result unequivocally shows that the G proteins bound to β2AR dimers, and not monomers, in the resting state are responsible for triggering the β2AR’s basal constitutive activity, which is sensitive to inverse agonism. It further suggests that the binding of the inverse agonists to β2AR induces conformational changes that decrease the β2AR dimer affinities for G proteins (**Fig. 2a**) and at the same time, hide the binding site(s) for G proteins in the dimer conformations.

Although the function of the GPCR dimers is to maintain the basal constitutive activity, it is critically important to realise that the GPCR dimers are metastable with lifetimes on the order of 0.1 s. Therefore, the GPCR dimers are dispersing continually, forming new dimers with other partner molecules all the time. In cells, this allows the basal constitutive activities of GPCRs to be maintained at certain levels, and yet the levels of the basal GPCR signals (the average number density of GPCR dimers at any time) are regulated globally, by modulating the GPCR number density by varying the expression levels of the GPCR, and locally, by temporarily and spatially varying the actin-based membrane skeleton and the membrane curvature^5,20–22^. In addition, the metastable GPCR dimers might play important roles in the activation of G proteins after the agonist addition.

The present results suggest that the major functions of some of the orphan GPCRs might be performed by the basal activities induced by their homo-dimers, as the GPCR family includes many orphan receptors^3,23^. Based on the results described in this report, we strongly suggest the pursuit of drug development strategies that emphasise the generation of more inverse agonists for various GPCRs, as well as drugs that enhance or suppress GPCR homo-dimer formation.

## Online content

Any methods, additional references, Nature Research reporting summaries, source data, extended data, supplementary information, acknowledgements, peer review information; details of author contributions and competing interests; and statements of data and code availability are available at http://doi.org/????????.

## Acknowledgements

We thank Prof. Hajime Fujisawa of Nagoya University for providing mouse L cells, Prof. Tatsuya Haga of Gakushuin University for providing the cDNAs encoding β2AR and Gα, Prof. Tobias Meyer of Stanford University for providing the cDNA encoding YFP-FKBP through Addgene (plasmid 201751), Prof. Michiyuki Matsuda of Kyoto University for the excellent suggestion about the method for establishing cell lines stably expressing target proteins at high levels, Assoc. Prof. Akira Hattori for the use of the PerkinElmer time-resolved fluorescence resonance energy transfer reader, EnVision Xcite, for the cAMP measurement, Drs. Taka-Aki Tsunoyama of OIST, Ko Hirosawa of Gifu University, Prof. Kenichi Suzuki of Gifu University, and all of the members of the Kusumi laboratory for valuable discussions, suggestions, and support, and Mr. Koji Kanemasa of incomings for assistance with figure preparation. This work was supported by Grants-in-Aid for Scientific Research from the Japan Society for the Promotion of Science (Kiban C to R.S.K. [17K07333], Wakate B to R.S.K. [26870292], Kiban S to R.S.K. [Co-PI, 18H05269], and Kiban B to R.S.K. [Co-PI, 17H03666], Kiban B to T.K.F. [16H04775], and Kiban S to A.K. [16H06386]), Grants-in-Aid for Innovative Areas (Molecular Engine) from the Ministry of Education, Culture, Sports, Science, and Technology (MEXT) of Japan to R.S.K. [18H05424], and the Joint Research Program of Biosignal Research Center of Kobe University to R.S.K. [301002]. WPI-iCeMS of Kyoto University is supported by the World Premier Research Center Initiative (WPI) of the MEXT.

## Author contributions

R.S.K. performed the single fluorescent-molecule tracking experiments and biochemical experiments. R.S.K., T.K.F., and A.K. developed and built the single-molecule imaging station. T.K.F. developed the analysis software. R.S.K. and A.K. conceived and formulated the project, evaluated and discussed the data, and wrote the manuscript, and all authors participated in revising the manuscript.

## Methods

### Cell culture, cDNA construction and expression in L cells, and treatment of cells with agonists and inverse agonists

L cells (a kind gift from H. Fujisawa of Nagoya University), which do not express β2AR^24^, were confirmed to be free of mycoplasma contamination using MycoAlert (Lonza), and were routinely cultured in Dulbecco’s Minimum Essential Medium (DMEM; Sigma-Aldrich) supplemented with 10% (v/v) FBS (Sigma-Aldrich), 100 units/ml penicillin (Wako), and 0.1 mg/ml streptomycin (Wako). Human β2AR was fused at its N-terminus to the ACP- or Halo7-tag protein (NEB and Promega, respectively) at the cDNA level, and cloned into the expression plasmid pEGFP-N1 (Clontech) after the removal of the cDNA region encoding EGFP. The ACP- and Halo7-tag proteins do not induce dimerisation^11,25^. For the genetic conjugation of mGFP to Gαs, the mGFP cDNA was inserted between the region encoding the 71st and 82nd amino acids of mouse Gαs (Origene) in the pcDNA3.1 (+) vector (Invitrogen)^26^, and for that to Gα, mGFP was fused to the N terminus of bovine Gα1 (a kind gift from T. Haga of Gakushuin University) ^27^ in the pEGFP-C1 vector (Clontech), after the removal of the cDNA region encoding EGFP. The details of the constructed cDNAs are shown in **Extended Data Fig. 1**. The L cells were transfected with these cDNAs using LipofectAMINE LTX (Life Technologies), according to the manufacturer’s recommendations. Transfected cells were seeded in glass-base dishes (35 mmϕ with a glass window of 12 mmϕ, 0.12~0.17-mm-thick glass; Iwaki), and cultured in Ham’s F12 medium (Sigma-Aldrich), supplemented with 10 % (v/v) FBS (Sigma-Aldrich), 100 units/ml penicillin (Wako), and 0.1 mg/ml streptomycin (Wako), for 24~48 h before microscope observations. The F12 medium was employed because it has a lower fluorescence background as compared with DMEM, which was used for routine culturing.

The agonist, isoproterenol (Sigma-Aldric), and the inverse agonists, ICI-118,551 (Tocris) and timolol (Wako), were dissolved in Hank’s balanced salt solution (Nissui) buffered with 2 mM piperazine-N,N’-bis(ethanesulfonic acid) (PIPES, Dojindo) at pH 7.4 (P-HBSS), at concentrations of 20, 200, and 200 μM, respectively. For the additions to the cells, 1 ml solutions of these reagents were added to the cells in 1 ml of the Ham’s F12 medium described above, providing the final concentrations of 10 μM isoproterenol, 100 μM ICI-118,551, and 100 μM timolol.

### Labelling of β2AR with the fluorescent dye ATTO594

ATTO594-Coenzyme A was custom-synthesised by Shinsei Kagaku, by conjugating ATTO594-maleimide (ATTO-TECH) to Coenzyme A-SH (New England Biolabs), followed by HPLC purification. The chemical structure of ATTO594-Coenzyme A was confirmed by liquid chromatography-mass spectrometry (LC-MS; LCMS-2010A EV, Shimadzu). The ACP-tagged β2AR expressed in the PM of the L cells was labelled with ATTO594, by incubating the cells with 2 μM ATTO594-Coenzyme A and 1 μM phosphopantetheine transferase (New England Biolabs) in Ham’s F12 medium without FBS at ~25°C for 20 min. These conditions have been known to achieve ~95% labelling of ACP^25^. The cells were washed with P-HBSS three times, and then subjected to microscope observations. The cells exhibiting the presence of typically ~1 ATTO594-ACP-B2AR spot/μm^2^ in the PM were used for single-molecule imaging experiments.

### Single fluorescent-molecule video imaging of live cells

The fluorescently-labelled ACP-β2AR expressed in the bottom PM was observed at the level of single molecules at 37°C, using home-built objective-lens-type total internal reflection fluorescence microscopes based on Olympus IX-70, Olympus IX-81, or Nikon TE-2000 inverted microscopes^4,28–30^, with a 100x objective lens, as optimised for the present research. Fluorescence images were projected onto a two-stage microchannel plate intensifier (C8600-03, Hamamatsu Photonics), coupled to an electron bombardment CCD camera (C7190-23; Hamamatsu Photonics) or an sCMOS camera (ORCA-Flash 4.0; Hamamatsu Photonics) operated at 30 Hz. For observations at a frame rate of 250 Hz, a CMOS camera (FASTCAM 1024PCI-II, Photron) was employed^31^. Fluorescent spots were identified by using an in-house computer program, as described previously^4^. The positions (x and y coordinates) of all of the observed single fluorescent-molecules were determined by a computer program that employed the method developed previously^32,33^. For the simultaneous dual-color single-molecule imaging, the images were spectrally split into two emission arms, each containing the same image intensifier and camera.

To maintain the cells at 37°C during the microscope observations, the entire microscope, except for the two detection arms and the excitation arm, was placed in a home-built microscope environment chamber made with thermo-insulating transparent vinyl sheets, and three heating circulators (SKH0-112-OT, Kokensya) were appropriately placed in the chamber to supply warmed air with slow circulation^30,34,35^. For further stabilisation, a plate heater (INUG2, Tokai Hit) was mounted on the microscope stage. The temperature of the surrounding air just above the stage was continually monitored during the observations with a digital thermometer (DT-312, Intermedical).

### Detection of colocalisation-codiffusion, and evaluations of β2AR dimer lifetimes and two-dimensional dissociation constants of β2AR dimers

The number densities of β2AR molecules existing as (true) dimers and the number density of the expressed β2AR molecules were determined by single-molecule imaging of ATTO594-ACP-β2AR^4^. Colocalisations of two fluorescent spots (molecules) of the same color (the same ATTO594 probe) or those of two different colors (YFP for BiFC and ATTO594) were detected when the two fluorescent molecules became localised within 220 nm of each other^4,30^. The number densities of incidental colocalisations at given molecular densities were subtracted^4^. The two-dimensional dissociation constant (2D-*K_D_*) of β2AR in the PM was determined by using the method previously developed by us^4^.

To directly detect the actual binding of two β2AR molecules, forming homo-dimers, we employed the bimolecular fluorescence complementation (BiFC) method^4,36^. The details of the BiFC method are described in the next subsection.

To determine the lifetime of β2AR dimers using single color observations (with ATTO594-ACP-β2AR), the duration of each colocalisation-codiffusion event was measured^4,11,29^, and after the durations of many events were determined, histograms showing the distribution of colocalisation durations were obtained. All of the histograms obtained from the image sequences obtained at a time resolution of 33 ms could be fitted by single exponential functions, which provided decay time constants (lifetime; *τ*_observed_). *τ*_observed_ was then corrected for the photobleaching lifetime of the fluorescent probe ATTO594 (*τ_bleach_* = 7.21 ± 0.81 s for 30 Hz [*n* = 300] and 0.812 ± 0.041 s for 250 Hz [*n* = 539]), to obtain the duration of apparent colocalisation (*τ_apparent colocalisation_*), using the equation [*τ_observed_*^-1^ - 2*τ_bleach_*^-1^]^-1^ ^4,11,29^. The effective dimer lifetime or the effective binding lifetime of proteins, *τ*, was evaluated by subtracting the duration of incidental colocalisation (without specific molecular interactions; *τ_incidental_*) from *τ_apparent colocalisation_*. For details, see our previous publications^4,29^. *τ_incidental_* was evaluated by using a monomer reference molecule, ACP linked to the transmembrane domain of the low-density lipoprotein receptor, called ACP-TM^29^, and was found to be 19 ± 0.46 ms (exponential lifetime; *n* = 241 examined dimers; 39 examined movies; No. of Freedoms = 97).

### Observations of the recruitment of Gαs and Gβγto βAR monomers and dimers

The recruitment of Gαs and Gβγ to βAR monomers and dimers was detected by observing the colocalisation-codiffusion of mGFP-conjugated Gαs and Gβγ with ATTO594-labelled β2AR monomers and dimers, at the level of single molecules. The colocalisation events lasting for longer than 66 ms (2 video frames) were analysed. The frequency of incidental colocalisation was estimated by superimposing an image sequence with one color on another image sequence with a different color that was artificially shifted by ~500 nm (horizontally), and was subtracted. The recruitment frequencies of Gαs and Gβγ to βAR monomers and dimers were normalised by the number densities of Gαs (or Gβγ) and β2AR protomers in the PM (per sec per μm^4^).

### The BiFC method for examining the formation of β2AR dimers (the interaction of two β2AR molecules at the molecular level)

The YFP-based BiFC(N) and BiFC(C) moieties employed previously^4,37^ were respectively fused to the C terminus of ACP-β2AR (**Extended Data Fig. 1**) in the pEGFP-N1 vector (Invitrogen; the cDNA region encoding EGFP was removed). These proteins were co-expressed in L cells, by transfection with LipofectAMINE LTX (Life Technologies) according to the manufacturer’s recommendations, and then ATTO594 was conjugated to the ACP moieties of ACP-β2AR-BiFC(N) and ACP-β2AR-BiFC(C). As the monomer reference molecule for the BiFC data, ACP-TM was employed and the YFP-based BiFC(N) and BiFC(C) moieties were fused to the C terminus of ACP-TM, as previously described^4^. Using dual-color single-molecule imaging (ATTO594 and mGFP), the number densities of the expressed molecules (the sum of the number densities of ACP-conjugated molecules) and those of the fluorescence spots of reconstituted BiFC molecules (spots/μm^2^) were determined.

### Evaluation of the changes of the cytosolic cAMP levels

The amounts of cAMP in the cytosol were measured by using the time-resolved fluorescence resonance energy transfer (TR-FRET) kit from Perkin Elmer (EnVision Xcite), according to the manufacturer’s recommendations. Since the L cells used for single-molecule experiments expressed very low levels of β2AR, and thus the basal level of cAMP in the resting state was below the detection limit, we first established a stable L-cell line expressing β2AR at high levels^38^, and used these cells to evaluate the changes of the cAMP levels in the cytosol before and after the additions of the agonist and inverse agonists. The drug treatments were performed as described at the end of the subsection, “Cell culture, cDNA construction and expression in L cells, and treatment of cells with agonists and inverse agonists”.

### Generation of artificial β2AR dimers and observations of Ca^2+^ mobilisation

To artificially generate stable homo-dimers of β2AR, we employed a chemical dimerising system based on the FK506-binding protein (FKBP)-ligand interface^39^. First, the cDNA encoding Halo7-β2AR-FKBP_F36V (β2AR-FKBP) was cloned into the expression plasmid vector pEGFP-N1 (Clontech) after the removal of the cDNA region encoding EGFP, and then the cDNA encoding FKBP_F36V was replaced by the cDNA encoding FKBP (obtained from the YFP-FKBP cDNA, the Addgene plasmid 20175 donated by T. Meyer^40^). Halo7-β2AR-FKBP expressed in L cells was fluorescently labelled by incubating the cells for 24 - 48 h with 5 nM Halo-tag conjugated with SaraFluor 650T (Goryo-Kayaku; SaraFluor650T-Halo7-β2AR-FKBP). The crosslinker for FKBP, AP20187 (Takara), was added at a final concentration of 5 nM during the image acquisition, and dimer formation was detected at the level of single molecules.

For Ca^2+^ mobilisation assays, a freshly-prepared 1 mg/ml DMSO solution of Fluo4-AM (Dojindo) was added to HBSS buffered with 10 mM (4-(2-hydroxyethyl)-1-piperazineethanesulfonic acid) at pH 7.4 (H-HBSS), containing 1.25 mM Probenecid (Dojindo) and 0.04 % (w/v) Pluronic F127 (Dojindo) (loading buffer), to obtain a final Fluo4-AM concentration of 5 μg/ml (4.6 μM), basically following the manufacturer’s recommendations. The L cells prepared as above for detecting dimers of SaraFluor650T-Halo7-β2AR-FKBP at the single-molecule level were incubated with the loading buffer at 37°C for 1 h and washed with H-HBSS three times, and then 500 μl H-HBSS was added for epi-fluorescence microscopic observations. Fluo4 fluorescence was recorded at 2 Hz. The stable dimers of SaraFluor650T-Halo7-β2AR-FKBP were induced by the addition of 5 nM AP20187, as described above. To determine the saturation levels of the fluorescence signal intensity of Fluo4 at higher concentrations of Ca^2+^, 1 μM (final concentration) ionomycin (Wako) was added to the cells to equalise the intracellular Ca^2+^concentrations with that outside the cells (1.3 mM).

### Software and statistical analysis

Superimpositions of image sequences obtained from dual-color single fluorescent molecule observations and single fluorescent spot tracking were performed using C++-based computer programs produced in-house, as described previously^30,32^. Statistical analysis was performed by the two-sided Welch’s t test (except for the data shown in **Extended Data Fig. 5c**, where the one-sided t test was used) or the Brunner-Munzel statistical test, by using the R Studio Software (R Foundation for Statistical Computing, https://www.R-project.org/). *P* values less than 0.05 were considered to be statistically significant. Linear and non-linear curve fittings were performed by using OriginPro 9.1 for Windows, with appropriate scripts.

### Data and code availability

All data generated or analysed for this study are available within the paper and its associated Extended Data Figures and Tables. All other data presented and software codes are available upon request from the corresponding author. A **Reporting Summary** for this paper is available.

