## Extended Data for "Metastable GPCR dimers trigger the basal signal by recruiting G-proteins"

### Extended Data Fig. 1

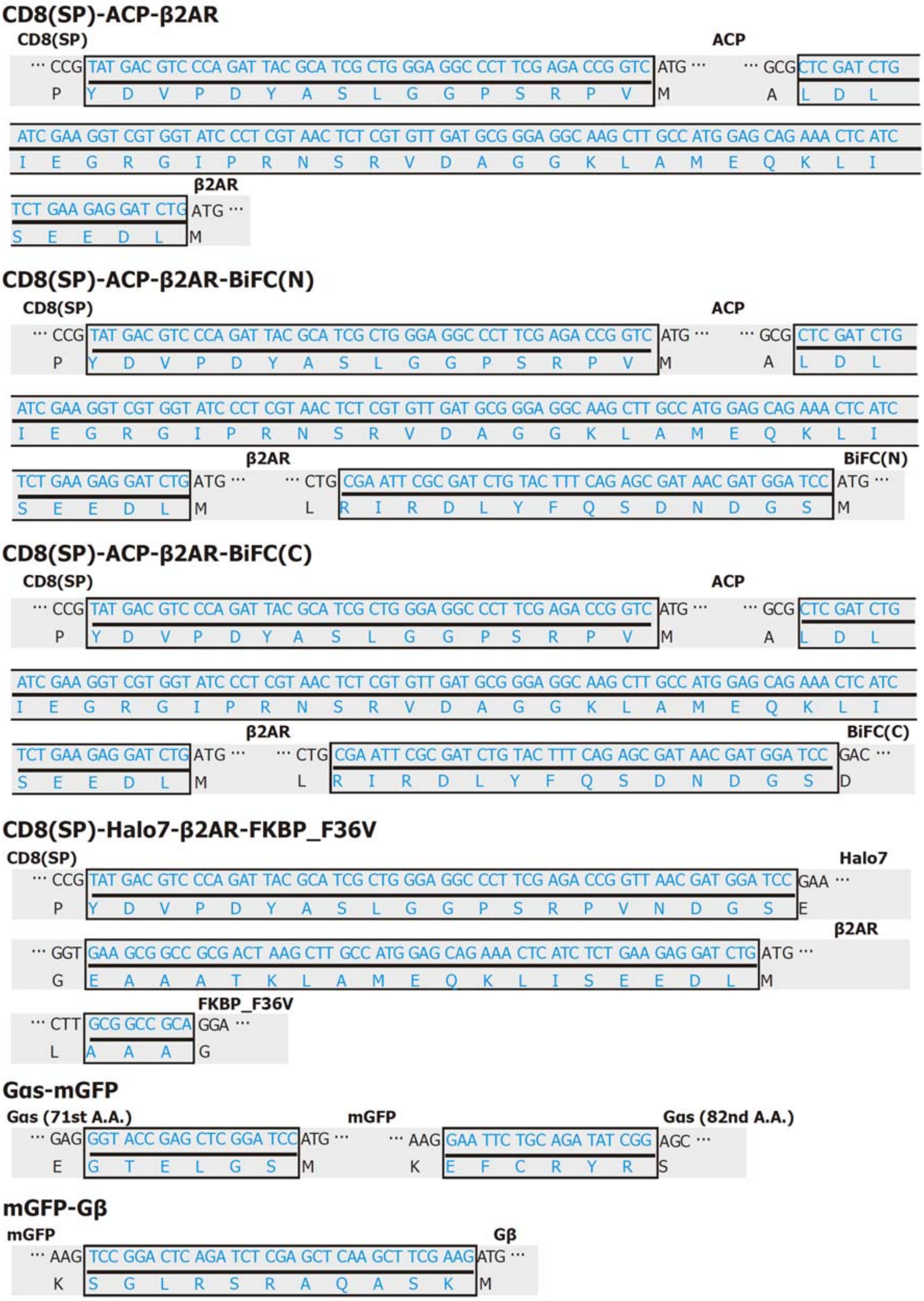


**Extended Data Fig. 1 | Schematic structures of the cDNA constructs used for protein expression in L cells, showing the added signal sequences (for β2AR), inserted tag proteins, linkers, and the target proteins.**

#### Extended Data Fig. 2

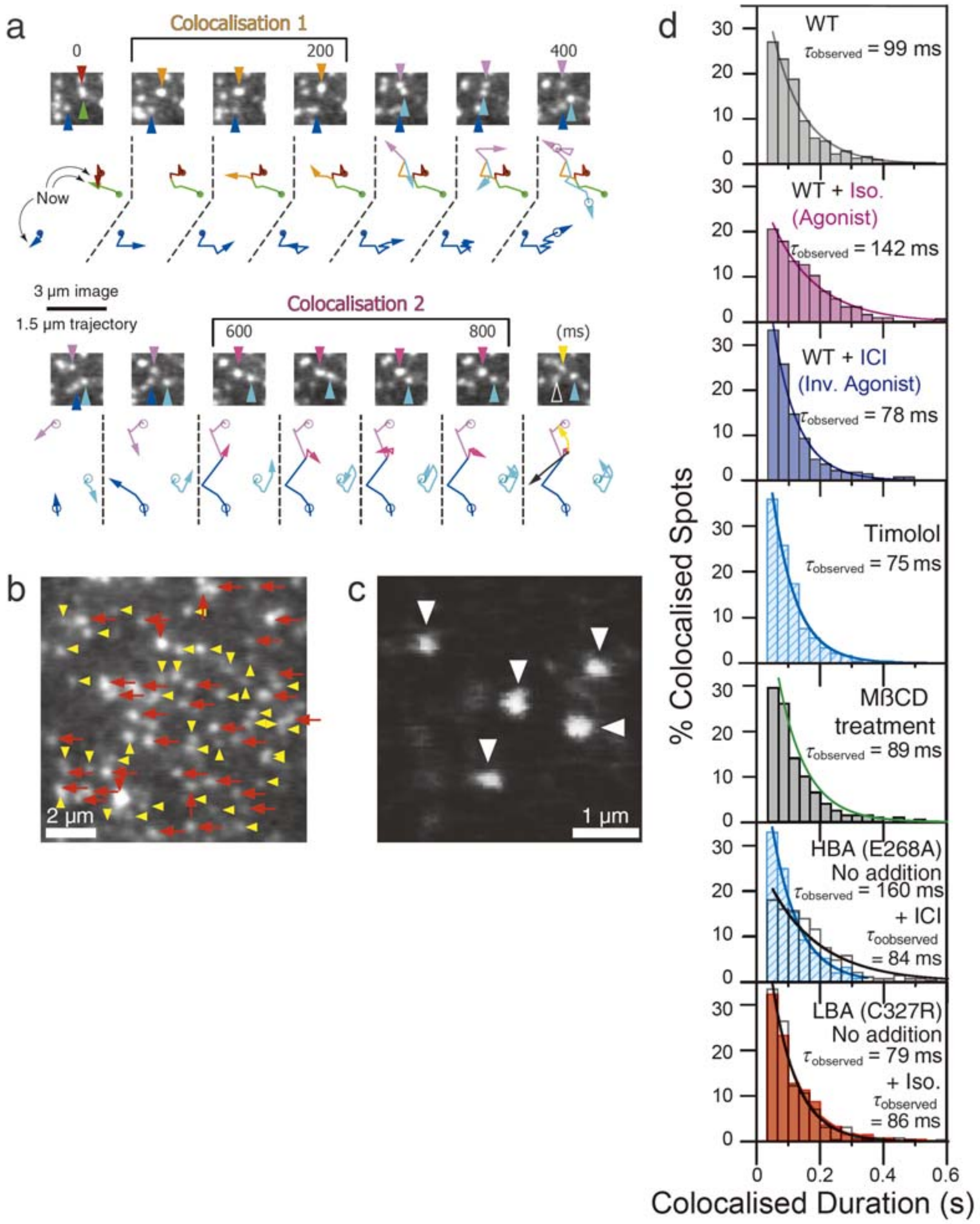


**Extended Data Fig. 2 | β2AR constitutively and continually forms transient homo-dimers with different partners in the PM (with lifetimes on the order of ~0.1 s), as confirmed by the BiFC method.**

**(a)** The observation duration is extended from that shown in **Fig. 1a**. After the first colocalisation lasting for 133 ms, the second colocalisation with another molecule started after 400 ms, and lasted for 200 ms.

**(b)** A representative image of single ATTO594-ACP-β2AR molecules, showing the co-existence of monomers (yellow arrowheads) and homo-dimers (red arrows) in the PM. Such images were used to obtain the graph shown in **Fig. 2a**, to evaluate 2D-$K_D$ of β2AR dimers.

**(c)** BiFC-YFP spots representing the reconstituted YFP in the dimer of ACP-β2AR-BiFC(N) and ACP-β2AR-BiFC(C).

**(d)** Histograms showing the distributions of the durations of colocalisation events (homo-dimer lifetimes) for WT β2AR as well as HBA- and LBA-mutants. Iso = isoproterenol (agonist). ICI = ICI-118,551 (inverse agonist). The exponential lifetime of colocalisation ($\tau_{observed}$) is shown in each box. $\tau_{observed}$ = the colocalisation lifetime without the correction for the photobleaching lifetime ($\tau_{bleach}$) and the subtraction of the incidental colocalisation lifetime ($\tau_{incidental}$) (see **Methods**). For the values after these corrections, see **Fig. 1c** and **Extended Data Table 1**.

### Extended Data Fig. 3

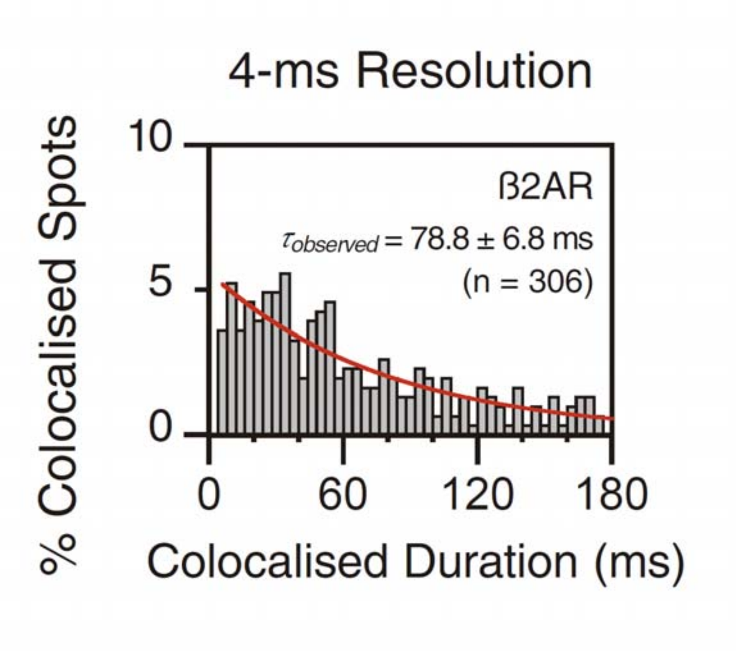


**Extended Data Fig. 3 | An observation frame rate of every 33 ms (normal video rate) is sufficiently fast for evaluating homo-dimer lifetimes of β2AR.**

The basic data for the second bar-graph (from the top) in **Fig. 1c**. The histogram shows the distribution of dimer lifetimes of β2AR obtained from observations made at a frame rate of every 4 ms (250 Hz, 8.3x enhanced from a normal video rate of 30 Hz; the red curve represents the best-fit single exponential function). The histogram could be fitted by a single exponential function, which provided a dimer lifetime of 78.8 ± 6.8 ms, before the corrections for photobleaching and incidental colocalisations (see **Methods**; $n$ = 306 examined dimers; 26 examined movies; No. of Freedoms = 57). After the corrections for photobleaching and incidental colocalisations (see **Methods**: "Detection of colocalisation-codiffusion, and evaluations of β2AR dimer lifetimes and two-dimensional dissociation constants of β2AR dimers"), the dimer lifetime as measured at a 4-ms time resolution was 78.8 ± 10.7 ms. Since this value was statistically indistinguishable from that obtained from the observations made at video rate (83.2 ± 6.4 ms, after the corrections for photobleaching and incidental colocalisations; see **Fig. 1c** and **Extended Data Table 1**), this result indicates that the observation frame rate of every 33 ms (normal video rate) employed in the present research is sufficiently fast for evaluating the homo-dimer lifetime of β2AR.

### Extended Data Fig. 4

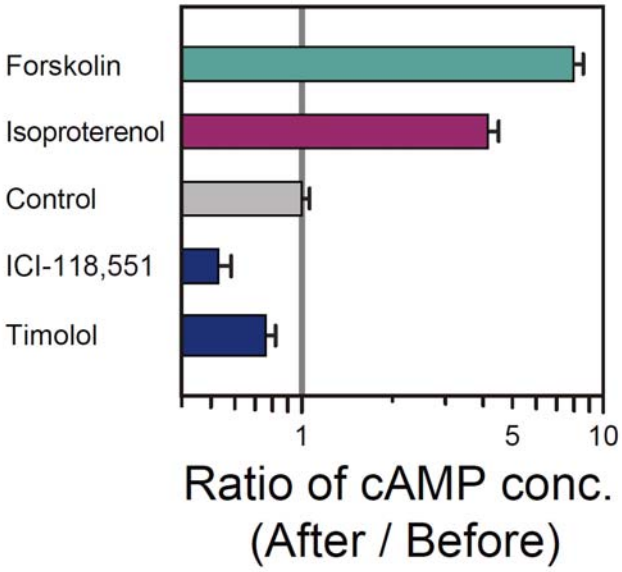


**Extended Data Fig. 4 | The basal constitutive activity of β2AR as evaluated by the cytosolic cAMP levels, using cloned L-cells stably expressing β2AR.** After the treatment with the inverse agonists, ICI-118,551 and timolol, the cAMP level was reduced, indicating that non-stimulated β2AR triggers the basal constitutive activity subjected to inverse agonism. The β2AR agonist, isoproterenol, increased the cAMP level. Forskolin, an adenylate cyclase activator, was used as a positive control.

#### Extended Data Fig. 5

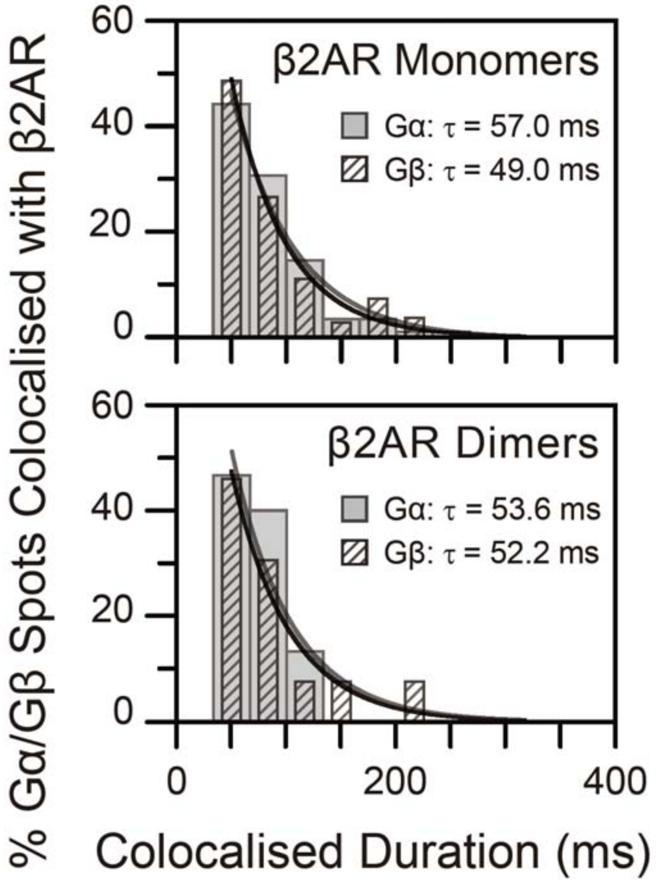


**Extended Data Fig. 5 | Histograms showing the distributions of the colocalisation durations of the trimeric G-protein (Gαs and Gβ) with β2AR monomers (top) and dimers (bottom). The maximal colocalisation lifetimes are on the order of 50 ms.** Considering the time resolution of 33 ms for the present observations, the colocalisation lifetimes cannot be determined correctly, and the values shown here are considered to be their maximal estimates.

### Extended Data Fig. 6

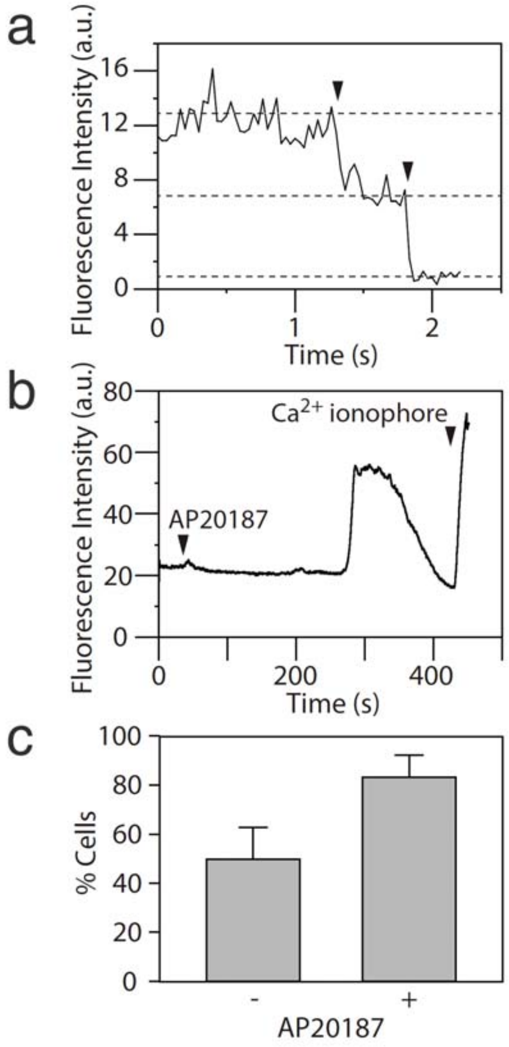


**Extended Data Fig. 6 | Artificially-induced β2AR dimers trigger the cytoplasmic signal without agonist application, as observed by $Ca^{2+}$ mobilisation, even when β2AR expression levels are as low as those used for single β2AR-molecule experiments.**

**(a)** β2AR dimers with lifetimes substantially longer than spontaneously-formed metastable dimers were induced by the addition of AP20187 to the cells expressing Halo-β2AR-FKBP (labelled with SaraFluor650-conjugated halo ligand; see **Methods**). Two-step photobleaching of the fluorescence spots was frequently observed after the addition of the FKBP crosslinker AP20187.

**(b)** The intracellular $Ca^{2+}$ concentration, monitored by the Fluo4 fluorescence intensity (per pixel), was increased (for durations less than 2 min) after the AP20187 addition, in larger fractions of the cells.

**(c)** AP20187-induced β2AR dimers enhanced the cytosolic $Ca^{2+}$ levels from the basal level ($P = 0.022$; $n = 18$ and 16 cells after the additions of AP20187 and H-HBSS [control], respectively). The cells that exhibited brief increases of the Fluo4 level higher than 7.5% of the saturation level (determined by the addition of the $Ca^{2+}$ ionophore, ionomycin) were counted.

**Extended Data Table 1** | Exponential lifetimes of β2AR homo-dimers (ms) with statistical parameters, after the correction for photobleaching lifetime and subtraction of the incidental colocalisation lifetime (19 ms; see **Methods**: "Detection of colocalisation-codiffusion, and evaluations of β2AR dimer lifetimes and two-dimensional dissociation constants of β2AR dimers"). The colocalisation events lasting longer than 66 ms (2 video frames) were analysed to avoid the influence of the shot noise.

| β2AR (WT, HBA, or LBA mutants) + added drugs | Dimer Lifetime Mean ± SEM (ms) | *P* value | No. of Examined Dimers | No. of Examined Movies | No. of Freedoms |
|---|---|---|---|---|---|
| WT | 83.2 ± 6.4[†, *1] | N. A. | 317 | 14 | 22 |
| WT + Isoproterenol | 128.9 ± 7.1[Y1] | 4.3 x $10^{-5}$ | 409 | 16 | 22 |
| WT + ICI-118,551 | 61.2 ± 3.1[Y1] | 0.040 | 280 | 20 | 23 |
| WT + Timolol | 57.2 ± 3.3[Y1] | 0.0018 | 238 | 15 | 17 |
| WT + MβCD (cholesterol depletion) | 72.0 ± 4.4[N1] | 0.42 | 200 | 7 | 27 |
| HBA | 148.2 ± 16.6[Y1, *2] | 2.4 x $10^{-6}$ | 297 | 17 | 17 |
| HBA + ICI-118,551 | 66.5 ± 4.6[Y2] | 4.0 x $10^{-9}$ | 255 | 15 | 8 |
| LBA | 61.2 ± 5.1[Y1, *3] | 0.035 | 254 | 14 | 17 |
| LBA + Isoproterenol | 68.7 ± 3.9[N3] | 0.40 | 276 | 16 | 17 |

[†]β2AR homo-dimer lifetime observed at a 4 ms resolution was 78.8 ± 10.7 ms (see **Extended Data Fig. 3**). The numbers of examined dimers and movies were 306 and 26, respectively, and the number of freedoms was 57.

*, [Y], and [N]. The results of the Brunner-Munzel tests. The distribution selected as the basis for the comparison is shown by the superscript, [*]. The three bases used here are indicated as *[1], *[2], and *[3]. The superscripts [Y] and [N] indicate that the distribution is or is not significantly different from the base distributions indicated by * (*P* values smaller or greater than 0.05), respectively.

**Extended Data Table 2** | Statistical parameters for 2D-$K_D$ of β2AR (WT) dimers (**Fig. 2a**), and for BiFC results (**Fig. 2b**).

**a**

| Addition of the Inverse Agonist | 2D-$K_D$ (copies/μm$^2$)<br>Mean ± SEM | No. of Examined Spots | No. of Examined Movies | No. of Freedoms |
|---|---|---|---|---|
| No addition | 1.6 ± 0.29 | 14,867 | 19 | 6 |
| + ICI-118,551 | 3.5 ± 1.1 | 9,101 | 12 | 3 |

**b**

| Examined Molecules | No. of Examined Spots | No. of Examined Movies |
|---|---|---|
| β2AR | 1,374 spots for ACP<br>397 spots for BiFC | 14 |
| ACP-TM | 363 spots for ACP<br>43 spots for BiFC | 8 |

**Extended Data Table 3** | The normalised frequencies of Gα and Gβγ recruitment to βAR monomers and dimers. The normalised frequencies of colocalisations = the number of colocalisation events per sec per μm$^4$ normalised by the number densities of Gαs (or Gβγ) and β2AR protomers. The frequencies of incidental colocalisations have been subtracted. See **Methods** "Observations of the recruitment of Gαs and Gβγ to βAR monomers and dimers".

| | | Monomers | *P* values against the resting state (monomers) | Dimer | *P* values against the resting state (dimers) | *n* (cells) |
|---|---|---|---|---|---|---|
| Gα | Resting | 1.6 ± 0.60 [*1] | N.A. | 1.7 ± 0.55 [*2, N1] | [N1] 0.92 (monomers vs dimers) | 22 |
| | +Isoproterenol (Agonist) | 3.6 ± 1.9 [N1] | 0.96 | 2.0 ± 2.6 [N2] | 0.49 | 16 |
| | +ICI-118,551 (Inverse Agonist) | 5.7 ± 2.7 [N1] | 0.056 | -0.058 ± 0.18 [Y2] | 0.012 | 12 |
| | +Timolol (Inverse Agonist) | -0.96 ± 3.9 [N1] | 0.66 | -0.035 ± 0.12 [Y2] | 0.0041 | 13 |
| | | Monomers | | Dimers | | *n* |
| | Resting | 0.26 ± 0.28 [*3] | N.A. | 0.32 ± 0.24 [*4, N3] | [N3] 0.83 (monomers vs dimers) | 9 |
| | +Isoproterenol (Agonist) | 0.17 ± 0.061 [N3] | 0.97 | 0.023 ± 0.015 [N4] | 0.14 | 23 |
| | +ICI-118,551 (Inverse Agonist) | 0.62 ± 0.43 [N3] | 0.34 | -0.072 ± 0.071 [Y4] | 0.025 | 7 |

**Extended Data Table 4** | The changes of the cAMP levels in the cytosol before and after the additions of the adenylate cyclase activator, forskolin, the agonist isoproterenol, and two inverse agonists, ICI-118,551 and timolol (see **Extended Data Fig. 4**).

| | cAMP amount normalized to that in control cells (Mean ± SEM) | *P* value against the control specimen* | *n* (wells) |
|---|---|---|---|
| + Forskolin (100 μM) | 7.97 ± 2.16 | $2.17 \times 10^{-7}$ | 12 |
| + Isoproterenol (10 μM) | 4.15 ± 0.35 | $1.59 \times 10^{-7}$ | 12 |
| Control | 1.00 ± 0.06 | N.A. | 24 |
| + ICI-118,551 (100 μM) | 0.53 ± 0.05 | $9.09 \times 10^{-7}$ | 18 |
| + Timolol (100 μM) | 0.76 ± 0.06 | $7.02 \times 10^{-3}$ | 19 |

*Using the two-sided Welch's t-test.

#### Supplementary Videos

**Supplementary Video 1. A Gαs molecule temporarily binds to a β2AR monomer.**

A representative movie showing that a single molecule of Gαs-mGFP (green) and a single molecule of ATTO594-ACP-β2AR monomer (red) diffuse in the PM and become colocalised for 0.1 s (yellow circle), in a resting L cell. The raw data for **Fig. 3a**. Replay = 45x slowed from real time. The horizontal width of the movie = 2 μm.

**Supplementary Video 2. A Gαs molecule temporarily binds to a transient β2AR dimer.**

A representative movie showing that, as soon as an ATTO594-ACP-β2AR dimer is formed from the two monomers (red), a single Gαs-mGFP molecule (green) binds to the β2AR dimer (yellow for 0.1 s), and then the complex is separated into two β2AR monomers and a single Gαs molecule. Observed in the PM of a resting L cell. Replay = 45x slowed from real time. The horizontal width of the movie = 2 μm.
